# Overexpression of *PpGL2* from *Prunus persica* enhanced soybean drought tolerance

**DOI:** 10.1101/2024.03.03.583192

**Authors:** Li Wei, Zhao Li, Li Dahong, Li Hong-yan

**Affiliations:** School of Biotechnology and Food Engineering, Huanghuai University, Zhumadian, Henan 463000, China; Garden Center, Huanghuai University, Zhumadian, Henan 463000, China

**Keywords:** Peach, PpGL2, Drought stress, Soybean

## Abstract

The HD-ZIP transcription factor family plays crucial roles in plant growth and abiotic stress responses. While its diverse functions and regulatory mechanisms are well-documented, its role in conferring abiotic stress tolerance in peaches remains largely unexplored. Here, we report the bioinformatics profile of PpGL2, a member of the HD-ZIP transcription factor family, and its integration into the soybean genome to assess its potential impact on drought tolerance. Localization studies in onion cells revealed nuclear localization of PpGL2-GFP fusion protein, while yeast hybridization experiments demonstrated its transactivation and DNA binding abilities. *PpGL2* overexpression under drought conditions led to reduced accumulation of reactive oxygen species and malondialdehyde compared to wild-type, decreased water loss rate, and increased chlorophyll content and relative water content. Additionally, *PpGL2* overexpression promoted plant height and root length under drought stress, accompanied by altered transcription levels of stress-related genes across different plant genotypes. Furthermore, *PpGL2* overexpression enhanced oxidative tolerance. Therefore, our findings suggest that *PpGL2* overexpression holds promise for enhancing soybean drought resistance, offering a novel approach to improving soybean drought resistance.

## 1 Introduction

Abiotic stress significantly affects plant growth, development, and crop yield, posing a substantial challenge to current agricultural economies. Among these stressors, drought stands out as particularly critical due to its frequent occurrence, prolonged duration, and wide-ranging impact. In response to stress signals such as drought, high salinity, and cold, plants activate specific functional genes through signal transduction processes[1]. Finally, these gene products prompt adaptive physiological and biochemical responses, facilitating plant survival[2]. For example, overexpression of *ATHB-6* enhances drought resistance in transgenic maize by activating reactive oxygen species-related genes[3]. Similarly, *PsnHDZ63* transgenic plants demonstrate improved phenotypes and physiological indicators under salt stress, suggesting enhanced salt stress tolerance due to increased PsnHDZ63 gene expression [4]. The overexpression of *Zmhdz4* and *Zmhdz10* genes in rice also enhances transgenic plant tolerance to drought and salt stress[5, 6]. Despite the identification and isolation of several stress-related transcription factor genes, their functions in soybean remain largely unexplored. Soybean (*Glycine max* (L.) Merrill.) is a nutritious legume widely utilized as a food source for humans and livestock due to its high protein and mineral content [7]. Moreover, soybeans are also a major oilseed crop, accounting for ∼59% of the world’s total oilseed production. However, like other crops, soybean growth and yield are significantly impacted by abiotic stressors such as drought and saline-alkali stress. For instance, drought alone can cause up to a 40% reduction in global soybean production [8], constraining both growth space and the potential of dryland as an alternative farmland. Various studies have demonstrated the pivotal role of plant homotypic leucine zippers (HD-ZIP) transcription factors in plant development and response to environmental stress, regulating tolerance through positive or negative regulation [9–11]. Therefore, the cloning of the peach tree HD-ZIP gene (*PpGL2*), analyzing its localization in onion surface cells and DNA-protein interaction sites, and *PpGL2* overexpression in soybean to assess its impact on drought resistance hold significant importance in identifying and harnessing high-efficiency stress resistance genes in plants.

HD-ZIP, a plant-specific transcription factor within the homeodomain (HD) transcription factor superfamily, is characterized by 60 (or 61) highly conserved homologous domains[12]. HD-ZIP proteins generally form homologous or heterodimeric complexes with DNA via the leucine zipper (LZ) domain, thereby modulating the expression of target genes[13]. Currently, HD-ZIP proteins have been identified in various plants including Arabidopsis[14], rice[15], sunflowers[16], tomatoes[17], and tea trees[18]. The classification of HD-ZIP transcription factors is comprehensive in Arabidopsis, where Ariel et al.[13] analyzed and identified 58 HD-ZIP proteins from the genome sequence. Based on their DNA binding specificity, physiological function, and structural domain characteristics, they were divided into four subfamilies, represented as HD-ZIP I-IV. Numerous plant HD-ZIP genes have been cloned and functionally characterized. For example, *AtHB6* and *AtHB7* genes in Arabidopsis are upregulated under water deficient conditions or upon external ABA application, indicating their crucial role in regulating the plant water deficit response[19]Similarly, yellow cauliflower exhibited four HD-ZIP proteins (CpHB4–CpHB7) responsive to water stress. Studies on the expression of four genes in yellow cauliflower reveal that CpHB4 and CpHB5 are downregulated by drought but unresponsive to ABA, whereas CpHB6 and CpPH7 are induced by both drought and ABA stress[20]. In rice, the HD-ZIP transcription factor OsHOX22 affects ABA biosynthesis and regulates drought and salt stress via ABA-mediated signaling pathways[21, 22]. Ariel et al.[23] found that salt stress induces the expression of HD-ZIP protein MtHB1 in the protoplast and lateral root meristem of *Medicago truncata*. MtHB1 exhibited an adaptive growth response to the environment, reducing root surface area exposed to severe salt stress[23]. Moreover, evidence from genetically modified plants suggests the involvement of HD-ZIP transcription factors in stress regulation. For instance, overexpression of the sunflower HD-ZIP protein encoded by HaHB-4 in Arabidopsis resulted in enhanced drought resistance and improved development[24]. Yan[25] introduced Zmhdz13 and Zmhdz14 into Arabidopsis, which conferred strong drought resistance and increased sensitivity to exogenous ABA. Various studies have reported instances of overexpressing transcription factors to bolster plant stress resistance, such as *OsMYB3R-2*[26], *CBF1*[27], *ABF2*[28], *bZIP17*[29], *BnbZIP2*[30], *OsbZIP46*[31, 32], and *OsTF1L* [15].

The HD-ZIP protein significantly influences plant growth and environmental stress response. Despite existing studies on the HD-ZIP protein, limited research investigates its impact on drought resistance in soybeans. This study cloned the *PpGL2* gene, encoding the HD-ZIP transcription factor, from the peach tree ‘Chunxi’. Subsequently, it conducted bioinformatics analysis and determined the subcellular localization of the PpGL2 transcription factor in the peach tree. Additionally, the studyoverexpressed the *PpGL2* gene in soybeans and study the drought resistance of transgenic soybeans. These findings aim to establish a theoretical foundation for further comprehensive investigations into the role of PpGL2 transcription factors in plant growth and development.

## 2 Materials and Methods

### 2.1 Test materials and processing

The test material comprises two-year-old potted seedlings of the peach tree cultivar “Chunxi” (*Prunus persica* (L.), grown in the greenhouse within the southern area of Huanghuai University. Robust young leaves were selected for RNA extraction and subsequent cDNA synthesis. The transient expression vector pCAMBIA1301(Fig S1)and *Escherichia coli* DH5α are both stored at the Garden Center Laboratory of Huanghuai University. The pMD18-T vector was procured from TaKaRa Bioengineering (Dalian) Co., Ltd. Reverse transcription, restriction endonuclease, PCR, and random primer kits were obtained from Bao Biotechnology (Dalian) Co., Ltd. DNA purification and recovery kits were acquired from Sagan Co., Ltd. (Shanghai). Antibiotics, plant growth regulators, etc., were sourced from Sigma Corporation in the United States, while all other reagents are of analytical grade, either imported or domestically produced. DNA polymerase I was purchased from Takara (Dalian, China). Key instruments and equipment include the PDS-1000 desktop gene gun (Bio Rad, USA) and a laser confocal microscope (ZEISS, Germany).

### 2.2 Primer design and synthesis

Total RNA was extracted from peach tree leaves using an RNA assay kit. Total RNA concentration and purity were measured using an ultra-trace nucleic acid protein analyzer, and RNA integrity was assessed using 1% formaldehyde denaturation gel electrophoresis. The method of cDNA synthesis involved the reverse transcription of 2 μg RNA with MMLV (H) reverse transcriptase to synthesize first-stranded cDNA. A 10 μL first chain reaction system was prepared, and DNA polymerase I was added to synthesize the second chain of DNA. Detailed synthesis steps were followed as per the instructions provided with the reverse transcriptase assay kit. Primer were designed for the NCBI peach *PpGL2* gene sequence (gene registration number: LOC18780448) using Primer Premier 5.0 software. The underlined sequences indicated corresponding enzyme cleavage sites (Table 1). The primers were synthesized by Shanghai Sagan Co., Ltd.

### 2.3 Quantitative PCR amplification

The quantitative RT-PCR mixture comprised 1 μL of cDNA synthesis reaction mixture, 5 μL of 1.2 μM primer premix, and 10 μ L SYBR premium ExTaq Perfect Real Time (TaKaRa Bio), adjusted to a final volume of 20 μL with water. Analysis was conducted using the DNA Engine Optic 2 system (Bio Rad). The PCR cycling conditions were: 95 °C for 10 s, followed by 20 s at 55 °C to 60 °C (depending on the gene), and 20 s at 72 °C, with a brief step at 78 °C for 2 s. This cycle was repeated 40 times. Fluorescence quantification at 78 °C was performed before and after incubation to monitor dimer formation. The *tubulin* gene served as internal reference standard. Additionally, a reaction mixture lacking reverse transcriptase was included to verify absence of genomic DNA contamination. Melting curve analysis and gel electrophoresis confirmed single DNA species amplification in all PCR experiments. Table S1 lists the primers utilized for real-time quantitative RT-PCR.

### 2.4 Bioinformatics analysis

The evolutionary analysis of PpGL2 transcription factors in peach trees is based on Ariel et al.’s (2007) HD-ZIP family classification method. From the Plant Transcription Factor Database (http://planttfdb.cbi.pku.edu.cn), 58 transcription factors were downloaded from the HD-ZIP family of Arabidopsis and compared with PpGL2 transcription factors in peach trees to construct a homologous evolution tree. Conservative domain prediction was conducted using the NCBI website (http://blast.ncbi.nlm.nih.gov/Blast.cgi) via BLAST; Multiple sequence alignment and hydrophilicity/hydrophobicity prediction were performed using DNAMAN 6.0 software. Physical and chemical properties were determined using the ProtParam online tool (http://web.expasy.org/protparam/) phylogenetic evolution tree generation and graphical reporting were achieved using Figtree software. Protein secondary structure prediction was completed on the SOPMA website (http://npsa-pbil.ibcp.fr/cgi-bin/npsa).

### 2.5 Recombinant plasmid construction

Using a gel recovery kit, the target band from PCR products was excised and cloned onto the pMD18-T vector for heat shock conversion of DH5α. Positive clones from the recipient cells were selected and sent for sequencing (Shanghai). The correct cloning vector was identified using restriction endonucleases BamHI and SacI for digestion, while the same endonucleases were employed for double digestion of the expression vector pCAMBIA1301 with GFP. Subsequently, the target gene and expression vector were combined at a ratio of 3:1 (50 ng:17 ng), followed by the addition of T4 ligase and overnight incubation at 4 °C. Following verification through transformation, enzyme digestion, sequencing, etc., the recombinant plasmid pCAMBIA1301-PpGL2-GFP was obtained. The fusion plasmid (35S: PpGL2-GFP) and control (35S: GFP) were introduced into onion epidermal cells via PDS-1000/He (Bio RAD, 1100 P.S.I.). Post-bombardment, the GFP signal was observed under a confocal microscope after a 36-h incubation in the dark at 24 °C on 1/2 MS solid medium.

### 2.6 Yeast hybridization

For transcriptional activation assays, the full-length ORF of PpGL2 fused with the GAL4 DNA binding domain in the PGBKT7 vector (Clontech, Japan) was transformed into the competent yeast strain Y2HGold. The pGBKT7 vector served as the negative control, while BD-p53 and AD-Sv40 large T antigen served as positive control. The transformed yeast cells were then introduced into Y2HGold cells containing *Aur1-C, Ade2, His3*, and *Mel1* reporter genes. These yeast cells were cultivated on Gal/AbA plates for four days under conditions of SD/Trp - and SD/Trp -/His -/Ade - /X-α-. To analyze DNA binding sites, the full-length ORF of *PpGL2* was fused with the GAL4 transcriptional activation domain in the pGADT7 vector (Clontech, Japan), and then co-transformed with the pHIS2 cis reporter vector (containing triple tandem CAAATTG repeat sequence) into yeast Y187 for determining DNA protein-protein interactions. Negative controls included p53 HIS2 and pGAD-PpGL2, while positive controls were p53 HIS2 and pGAD-Rec2-53. The pHIS2 mcis vector mirrors pHIS2 cis but features a binding sequence with four G bases (CGGGTTG). Co-cultured yeast cells were then grown on SD/Trp -/Leu -/His plates with or without 3-AT for 1 week.

### 2.7 Soybean planting and drought treatment

The experiment utilized the Williams 82 soybean variety. The genetic modification method should be consulted from literature [33]. First, soybean seeds were soaked in water for 6 h, then spread them evenly on moistened filter paper for germination. Once embryonic roots reach ∼1.0-1.5 cm, transfer them to quartz sand and irrigate with 1/2 Hoagland nutrient solution. Cultivate the seeds in a growth chamber set at 25 °C with a relative humidity of 60–70% and long days (16 h light/8 h dark). When the first pair of true leaves appear, select plants of equal height and subject them to 24 h of darkness for further growth. Plants were watered with 1/2 Hoagland nutrient solution containing 20% mannitol for drought treatment. Root and leaf samples were collected at 0 h, 3 h, 6 h, 12 h, and 24 h, then frozen with liquid nitrogen, and stored in an ultra-low temperature refrigerator. Three sets of biological replicates were included for each time point and tissue.

### 2.8 Measurement of antioxidant enzyme activity, malondialdehyde and chlorophyll concentration

Following the specified method, plants underwent a 2-week drought treatment. Antioxidant enzyme activity, including superoxide dismutase (SOD), peroxidase (POD), and catalase (CAT), as well as malondialdehyde (MDA) and chlorophyll content, was measured using the previously described method[34]. Three biological replicates were performed per treatment, and all experiments were repeated thrice.

### 2.9 Expression analysis of stress response genes

Soybean seeds were planted in vermiculite-filled pots. Upon emergence, seedlings were transferred to plastic containers with half-strength Hoagland solution (pH=6.5). One week post-transplantation, seedlings were treated with half-strength Hoagland solution supplemented with 15% mannitol. Root samples were collected 48 h post-treatment, immediately frozen in liquid nitrogen, and stored at −80 °C until analysis. Total RNA was isolated and reverse transcribed to obtain cDNA, serving as a template for real-time quantitative RT-PCR. Pressure-related gene expression (Table S1) was quantified from root samples.

Each stress treatment underwent three biological replicates. The mRNA level of the *Tubulin* gene served as a control for analysis.

### 2.10 Data Analysis

The data statistical analysis was completed using Excel 2013 and SPSS 17.0.

## 3. Results

### 3.1 Cloning of peach tree *PpGL2* gene and its overexpression in soybean

Using peach tree ‘Chunxi’ leaf cDNA as a template and employing PpGL2-F and PpGL2-R primers (Table S1), PCR product electrophoresis results reveals a clear, single target band ∼2200 bp long, consistent with expectations. Sequencing confirmed the *PpGL2* gene’s length in peach trees at 2253 bp, encoding 750 amino acids. Constructing a pCAMBIA-PPGL2 eukaryotic expression vector utilized a dual enzyme digestion and recombination method. Subsequent transformation of PPGL2 into soybean (T0 generation) was achieved via *Agrobacterium tumefaciens*. PCR amplification and electrophoresis using T1 soybean leaf genomic DNA as a template show clear 2200 bp bands in transgenic linesL1∼L7(Fig S2). Southern blot analysis showed positive reactions only in L1, L2, L5, L6 and L7. In contrast, wild-type soybean (WT) and false positive lines (L3 and L4) lack this band (Figure S1A). The five transgenic strains were screened to the T3 generation using hygromycin-containing medium, and L5, L6 and L7 have multiple copies, resulting in the identification of stable *PPGL2* gene inheritance in L1 and L2 transgenic soybean strains (Fig S2).

### 3.2 Bioinformatics analysis of PpGL2 transcription factors in peach trees

Figure S2 demonstrates that Arabidopsis harbors 58 transcription factors within the HD-ZIP family, categorized into HD-ZIP I-IV groups. Peach tree PpGL2 transcription factors exhibit a close relationship with Arabidopsis AT4G21750’s HD-ZIP IV transcription factors, suggesting PpGL2’s that PpGL2 belongs to the HD-ZIP IV subfamily of transcription factors in evolutionary terms.

The predicted conserved domain of PpGL2 in peach trees (Figure 1) reveals the START domain of the PpGL2 transcription factor spans amino acid sites 1 to 215, with homologous heterotypic domains (HD) ranging from 60 to 130, followed by a leucine zipper structure. Aligning the PpGL2 transcription factor of peach tree with walnut (*Juglans regia*, XP-018820351.1), *Jatropha curcas* (XP-012091395.1), *Quercus suber* (XP-023904570.1), apple (*Malus domestica*, XP-008384034.1), *Gossypium arborum* (KHG19735.1), cocoa (*Theobroma cacao*, XP-007026002.1), wild tobacco (*Nicotiana attenuate*, OIT00329.1), and grape (*Vitis vinifera*, XP:002266688.1), a multiple alignment of amino acid sequences of HD-ZIP transcription factors was conducted, including Arabidopsis AtPDF2 (AT4G04890.1) and AtML1 (AT4G21750.1) (Fig S3). Results indicate the highest homology with apple at 92.41%, whereas the lowest homology with AtPDF2 stands at 81.60% (Figure 1). Additionally, HD-ZIP transcription factors of these species all feature a START domain. BLAST homology search and alignment of the PpGL2 amino acid sequence in peach trees revealed the protein’s residue number as 750, with a relative molecular weight of 8.20 × 10^4^ Da and an isoelectric point of 5.87. Alkaline amino acids constitute approximately 12.7%, while acidic amino acids constitute ∼11.7%. Aromatic amino acids account for 5.9%.

**Figure 1:**
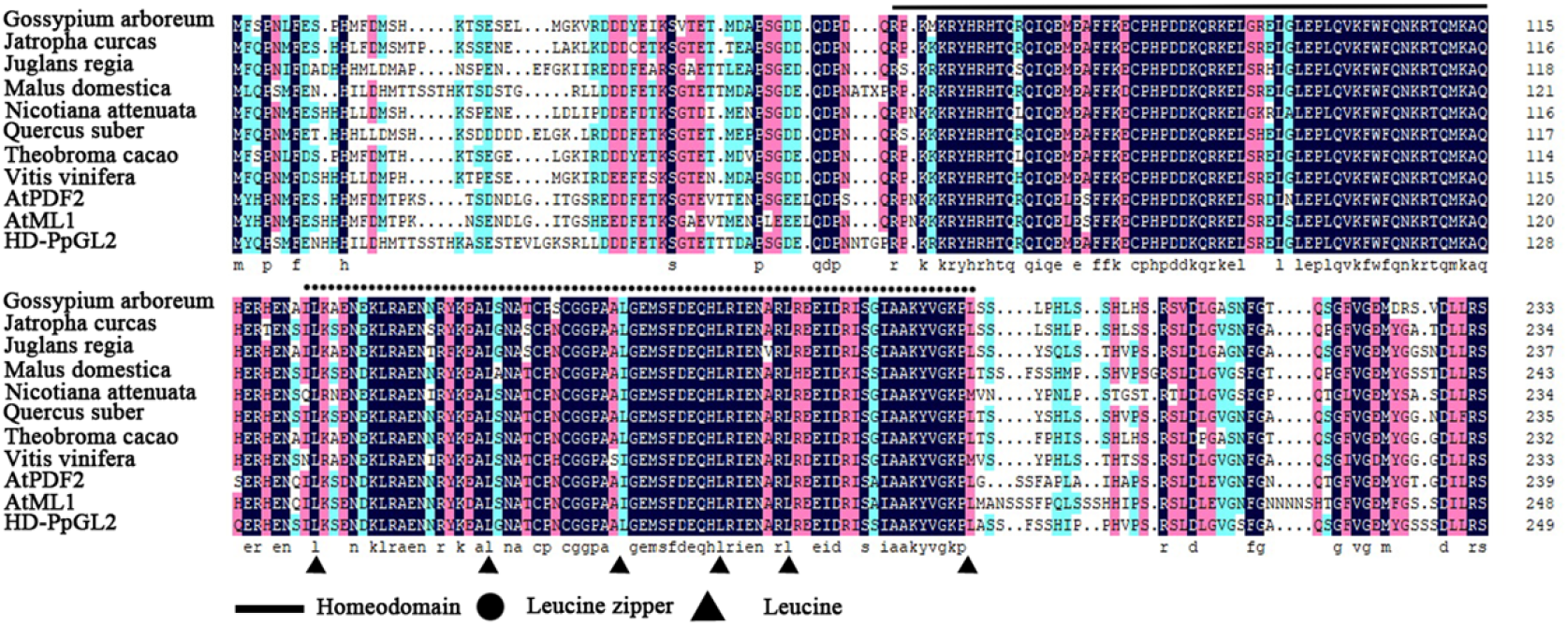
Sequence analysis of PpGL2

### 3.3 Peach tree PpGL2 transcription factor localization in the nucleus and activation of DNA

To analyze PpGL2 localization in cells, we constructed a PpGL2-GFP fusion vector (35S: PpGL2-GFP) and transformed into onion epidermal cells using a gene gun. Figure 2 illustrates that the GFP signal localized exclusively within the nucleus of onion cells in the PpGL2-GFP fusion vector (Figure 2-D), contrasting with the uniform distribution of GFP throughout the entire cell in the control (35S: GFP) (Figure 2-A). These findings affirm PpGL2’s nuclear localization. To investigate PpGL2 gene localization, we utilized the yeast single-hybridization method. The results showed robust growth of PGBKT7-PpGL2 and positive control strains on SD/Trp - medium, thriving in the presence of 300 μ Good growth was evident on SD/Trp -/His -/Ade - medium supplemented with g/L gold basidioside A (AbA), exhibiting α-Galactosidase activity. Conversely, the negative control only thrived on SD/Trp - medium, displaying no α-Galactosidase activity (Figure 3-A), indicating PpGL2’s transcriptional activity in yeast cells. To explore PpGL2’s DNA binding activity, we employed the yeast single-hybridization method. Figure 3-B demonstrates that only positive controls (pGAD-PpGL2 and PHIS2 cis) thrived on SD/Trp -/Leu -/His medium supplemented with 40 mmol · L-1 3-aminotriazole (3-AT), while negative controls did not grow. Furthermore, we constructed a pHIS2 mcis vector and co-transformed it with pGAD-PpGL2 into yeast cells. Figure 3-C illustrates that only the positive control grew on SD/Trp -/Leu -/His - medium containing 20mmol · L-1 3-AT, indicating PpGL2’s specific binding to the palindrome sequence CATTAAATG, thereby activating its reporter gene expression in yeast cells.

**Figure 2:**
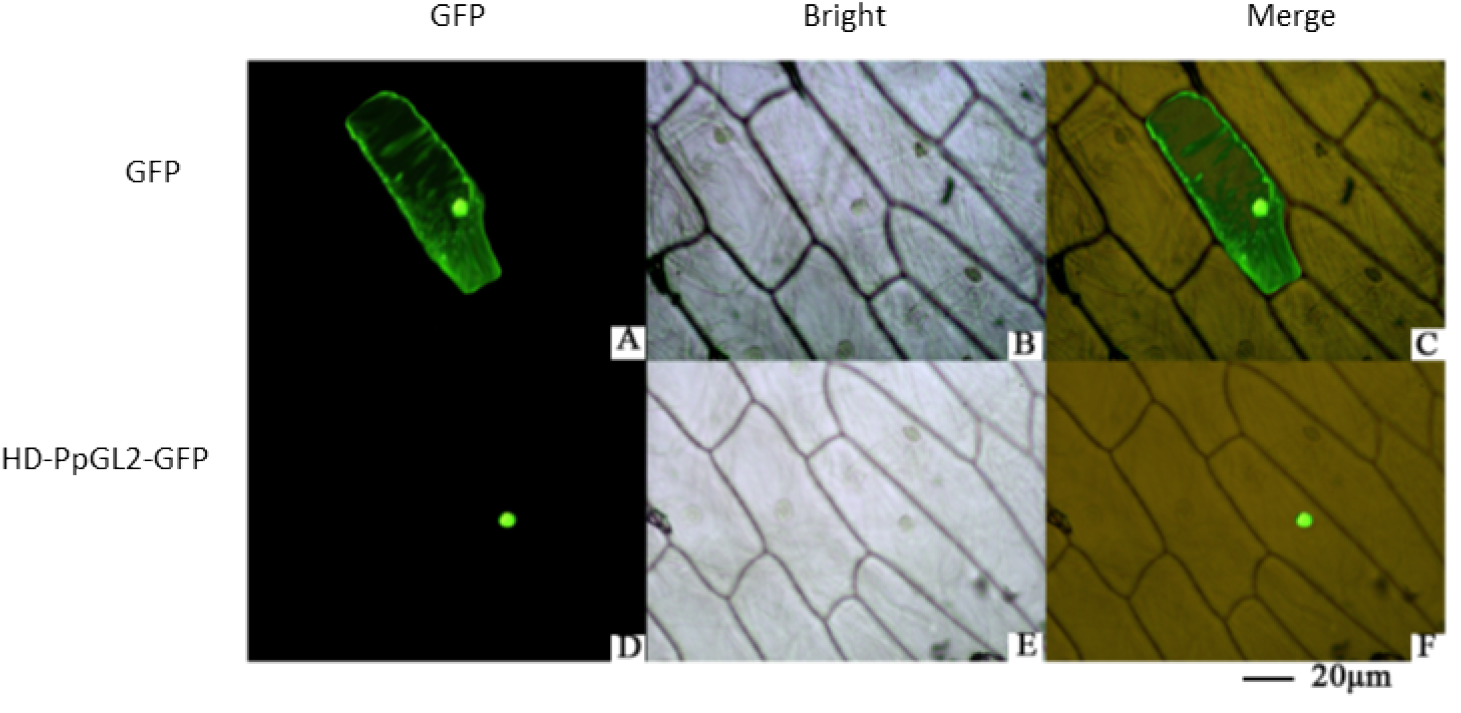
Subcellular localization of PpGL2 in onion epidermal cells

**Figure 3:**
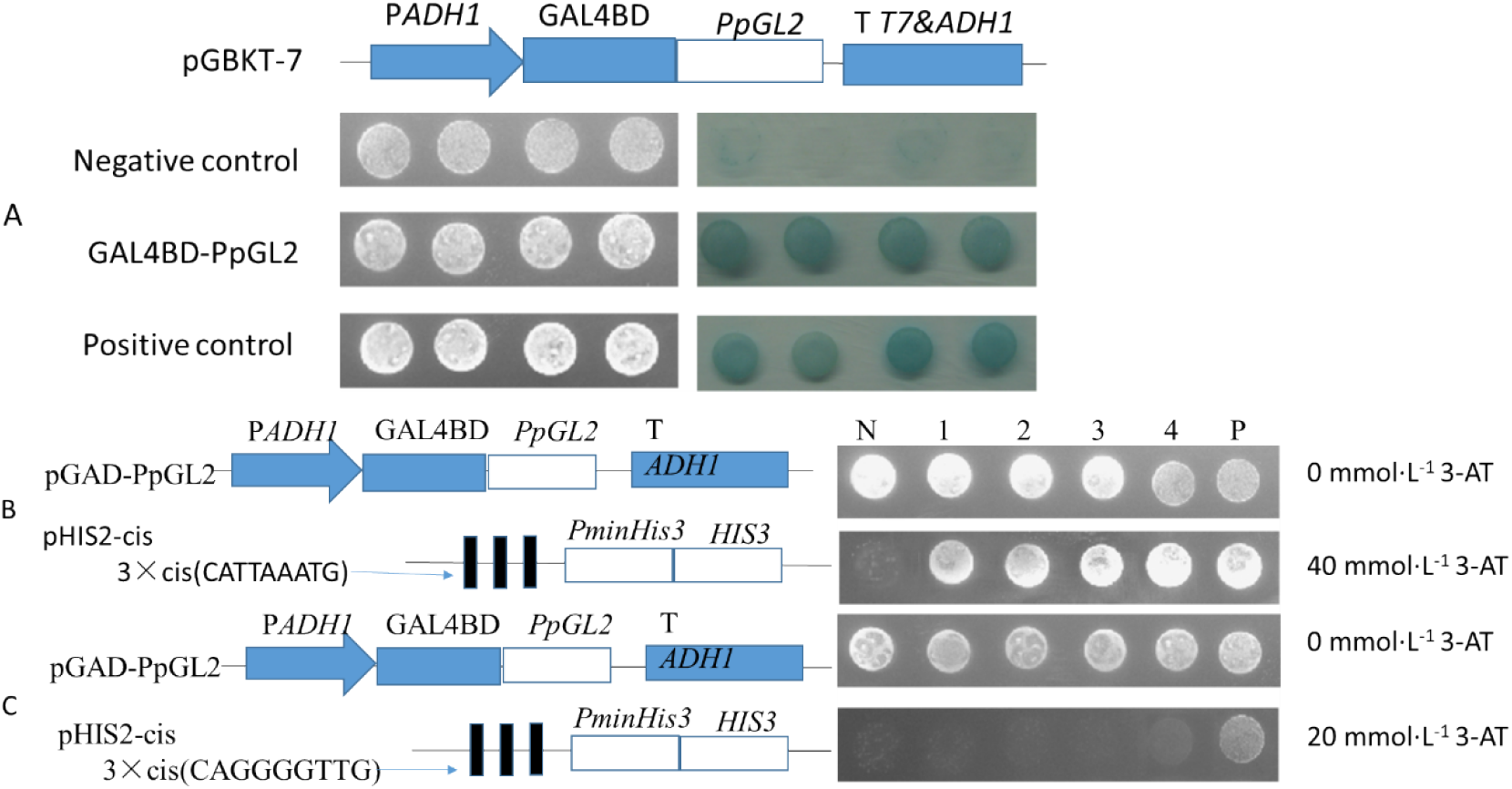
Transactivation activity and DNA-binding assays of PpGL2. A) Transactivation assay of PpGL2; B and C) DNA-binding assays of PpGL2. 1-4, with four different colonies containing positive transformants; N,negative control; P,positive control.

### 3.4 Overexpression of *PpGL2* enhances soybean drought resistance

Mannitol treatment was administered to transgenic strains. After a 10-day period, seedling survival rates were counted, and soybean plant height and root length were measured under stress conditions (Fig S4). Results indicated that stress treatment inhibited soybean root growth in both WT and overexpressing strains, albeit to varying extents. Notably, overexpressing strains exhibited significantly milder inhibition. In comparison to WT, overexpressing soybean strains displayed longer root length and greater plant height (Figures 4-B,C). Drought stress was applied to soybeans cultivated in nutrient-rich soil. Under normal conditions, no notable differences in growth status or phenotype were observed between WT and overexpressing strains. Following drought stress, both WT and overexpressed soybean lines exhibited leaf dehydration and plant wilting, although overexpression resulted in a lower dehydration rate compared to WT. Upon one week of rehydration treatment, overexpressing soybean strains commenced normal growth recovery, with most plants surviving. Conversely, the growth status of WT strains showed insignificant improvement, with most plants ultimately wilting and perishing (Figure4-D). Notably, the survival rate of overexpressed plants significantly surpassed that of WT(Figure4-E). Additionally, the chlorophyll content in overexpressing soybean plants before and after rehydration exceeded that of WT plants (Figure 4-A). These findings suggest that PPGL2 enhances the drought stress tolerance of transgenic soybeans.

**Figure 4:**
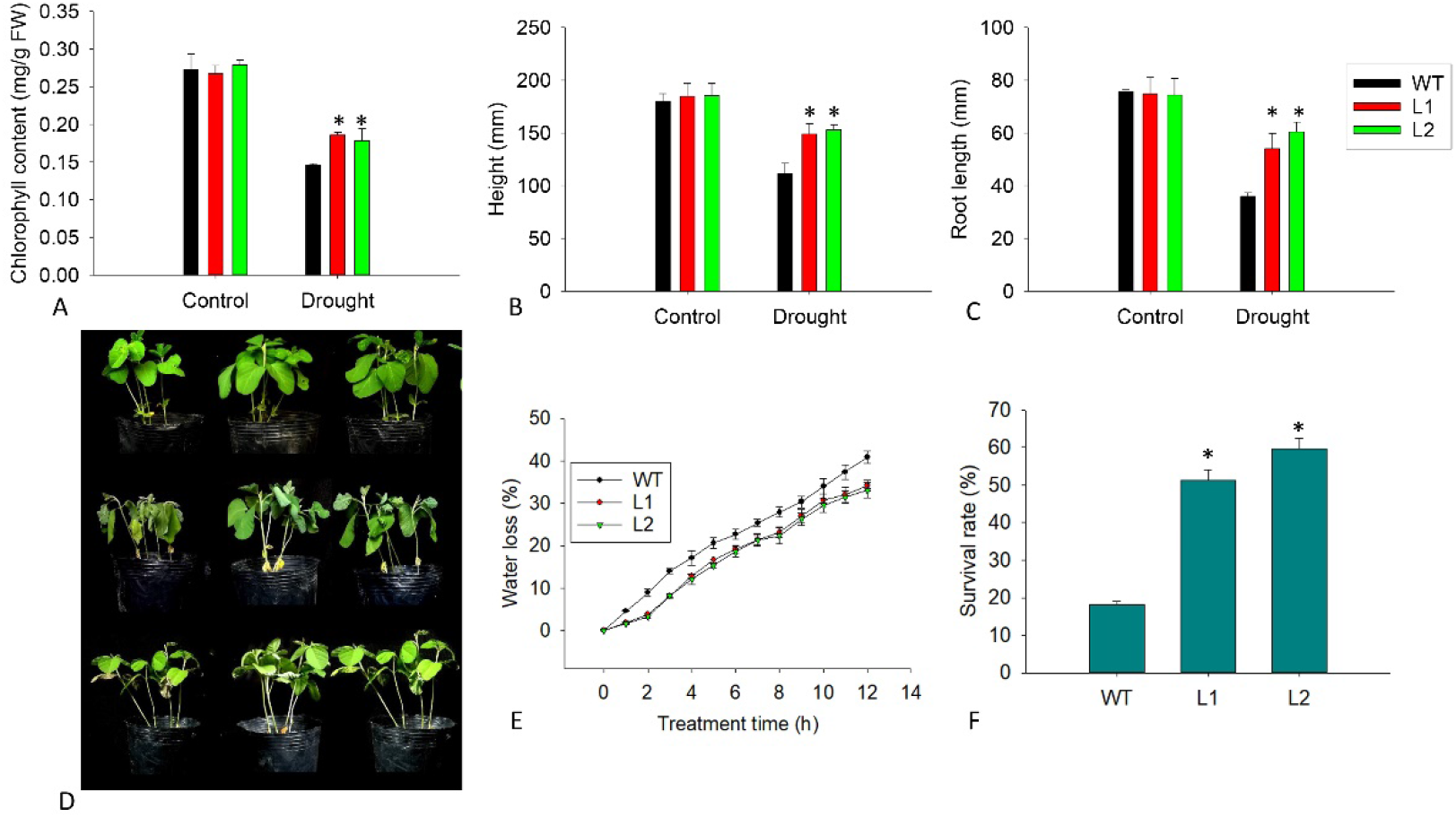
Enhanced drought tolerance in transgenic soybean. (A) Chlorophyll content during drought treatment of 35-day-old transgenic plants in a greenhouse for five days and then recovery for five days. Bar = 8 cm. (B), height (C) root length. (D) after recovery from drought stress. Data represents mean ± SD (∗p < 0.05). (E) water loss (F) Survival rate after 5-day drought treatment and four days of recovery. Values are mean ± SD (n = 50, ∗P < 0.05).

### 2.5 PPGL2 enhances the antioxidant capacity of genetically modified soybeans under drought stress

The RWC of overexpression plants is significantly higher than that of the control under drought stress (Figure5-A). Drought stress induces severe oxidative damage in plants, leading to significant accumulation of malondialdehyde (MDA) and increased electrical conductivity in soybean leaves. However, overexpressing strains exhibit significantly lower MDA and electrical conductivity levels compared to WT (Figure 5-B and C). The NBT (blue) and DAB (brown) staining results showed fewer blue and brown leaves in overexpressing strains under drought stress, indicating reduced accumulation of H_2_O_2_ and O_2_ in leaves (Fig S5). Antioxidant enzyme activity analysis demonstrates increased SOD, CAT, and POD activities in WT strains post-stress; yet, overexpressing strains exhibited higher activities (Figure 6-D-F). These findings suggest that *PpGL2* reduces reactive oxygen species (ROS) accumulation and mitigates oxidative damage by increasing antioxidant enzyme activity.

**Figure 5:**
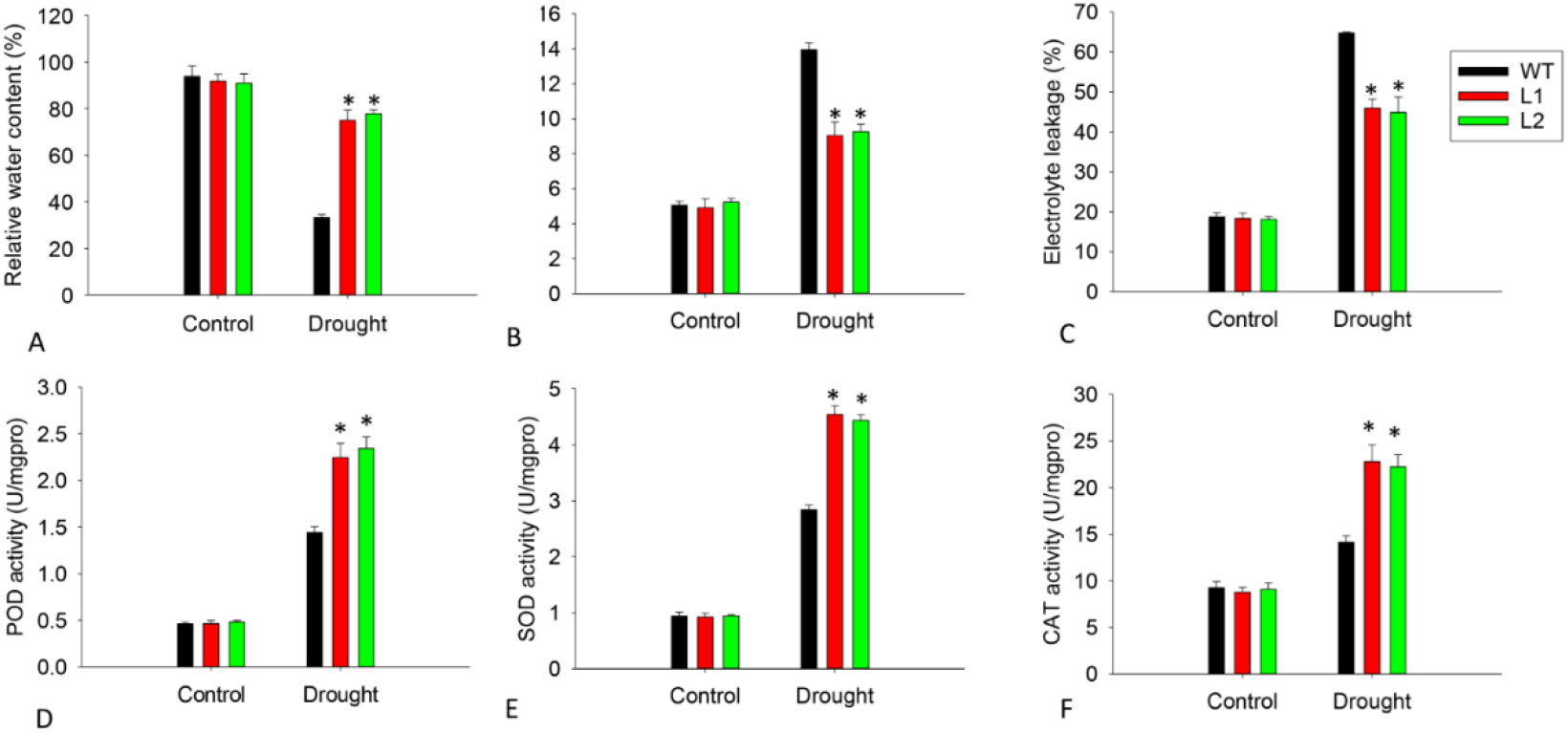
Overexpression of *PpGL2* decreased ROS production and oxidative damage under drought stress. (A) Relative water content. (B) MDA content. (C) electrical conductivity. (D-F) POD, SOD, and CAT activities. Error bars represent ± SD (n = 3). Asterisks denote significant differences (as compared with the control group): * p < 0.05.

**Figure 6:**
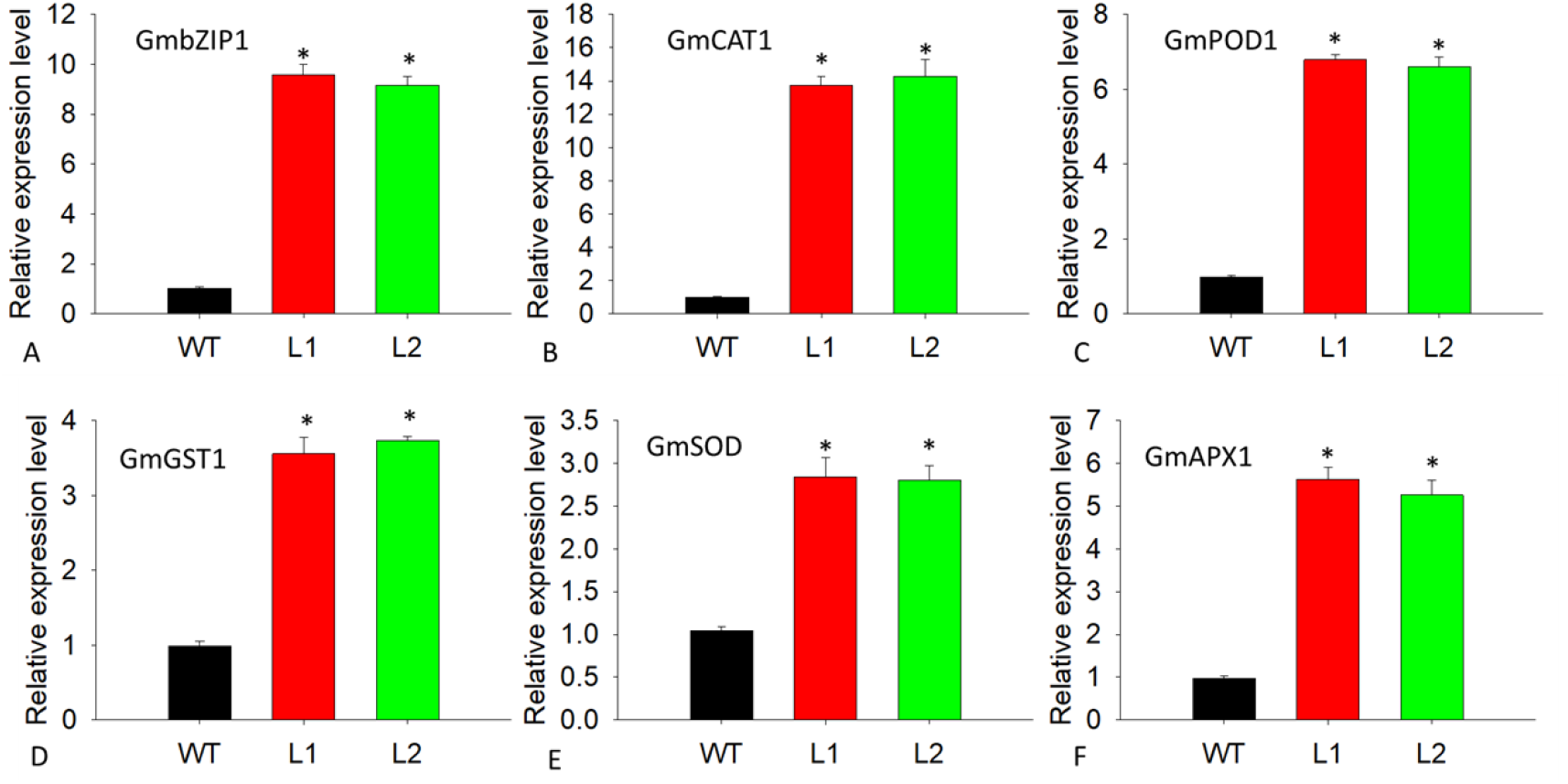
Relative expression levels of stress/ABA-responsive genes in transgenic Arabidopsis plants under drought stress

### 2.6 PPGL2 promotes the expression of stress response related genes in transgenic soybeans

The transcription levels of stress-responsive genes in WT and overexpressing soybean strains under drought stress were analyzed. Overexpressing strains show significantly higher expression levels of antioxidant enzyme-related genes (*GmSOD, GmPOD1, GmAPX1, GmGST1, GmCAT1*) and transcription factor *bZIP1* compared to WT strains (Figure 6). This underscores PpGL2’s role in promoting the expression of antioxidant enzymes and stress-responsive genes, thereby enhancing the drought resistance of transgenic soybeans.

## 3 Discussion

The HD-ZIP protein, unique to plants in the homologous box protein family, contains a highly conserved homologous heterotypic domain (HD) connected to an LZ domain at the carboxyl terminal. It binds to target DNA as dimers through the LZ domain[13]. This study reveals a homologous heterotypic domain (HD) at the N-terminus of the PpGL2 transcription factor, followed by a leucine zipper structure (Figure 2). Numerous studies confirmed the nuclear localization of HD-ZIP proteins [3, 35]. Similarly, our study demonstrates the nuclear localization of the PpGL2 transcription factor in onion epidermal cells through subcellular localization (Figure 4). Previous research indicates the ability of HD-ZIP IV protein to bind to the palindrome sequence CATT (A/T) AATG[13]. Our study, employing yeast single hybridization, illustrates the transcriptional activation of the *PpGL2* transcription factor in yeast cells. The DNA binding site is the CATT (A/T) AATG palindrome sequence (Figure 5). The Arabidopsis genome harbors 58 members of HD-ZIP, classified into four subtypes (I-IV) based on phylogenetic relationships. Arabidopsis HD-ZIP transcription factors, *AtPDF2* and *AtML1*, regulate Arabidopsis trichome differentiation and development. They also influence plant growth and development through a plant hormone signaling network[36]. Conservative sequence and domain analysis in our study suggest high conservation and homology of *PpGL2* with *AtPDF2* and *AtML1*. This implies that *PpGL2* may have similar functions. Studies show that Arabidopsis plants overexpressing the maize *ZmHDZ10* transcription factor gene exhibit enhanced tolerance to drought and salt stress[37]. Similarly, overexpression of *PpGL2* in soybean confers stronger drought resistance than wild-type plants, supported by morphological, physiological, and molecular evidence. Non-biological stress expression analysis reveals dehydration treatment-induced transcriptional upregulation of *PpGL2*. Mannitol treatment exhibited lesser impact on the height and main root elongation of transgenic seedlings compared to the wild type. Net photosynthesis decreases post-drought stress. The complex drought resistance mechanism involves different plant development stages, involving cells, tissues, organs, and physiological and biochemical processes[38]. When subjected to drought stress, 35-day-old soybean plants exhibit more severe damage symptoms (wilting, yellowing, and necrosis) in WT compared to overexpressing strains. Compared with WT, the relative water content (RWC) and water loss rate post-drought stress are two typical physiological parameters used to study the relative tolerance and resistance of crops under drought stress conditions[39]. Overexpressing transgenic plants demonstrate higher RWC and lower water loss rate post-drought stress. Oxidative stress activation under various stresses may lead to ROS increase (such as H_2_O_2_)[40]. MDA serves as a marker for oxidative stress-induced damage. Post-drought treatment, transgenic plants showed significantly reduced H_2_O_2_ and MDA levels compared to wild-type plants. Moreover, SOD, POD, and CAT content significantly increased in overexpressing strains, while only a slight increase was observed in WT. Overexpressed plants may accumulate more antioxidant enzymes under drought stress to resist ROS and repair oxidative damage. Additionally, stress-related gene transcription levels (*bZIP1, GmSOD, GmPOD1, GmAPX1, GmGST1, GmCAT1*) are higher in *PpGL2* overexpressing strains compared to the wild type. These findings suggest enhanced soybean drought tolerance during the nutritional growth stage upon PpGL2 overexpression.

In summary, our study investigated the bioinformatics of peach transcription factor PpGL2 and its cellular localization. Phylogenetic analysis highlights the evolutionary conservatism of peach PpGL2. Under drought stress, transgenic plants overexpressing PpGL2 exhibit elevated SOD, POD, and CAT activities, and increased soluble protein accumulation in plant cells, while MDA and electrical conductivity increased compared to controls. This enhances phenotype and resistance. Overexpression of soybean under drought stress resulted in higher chlorophyll content, plant height, and root length compared to controls. Our survey underscores the conservation of homologous evolution of peach HD-ZIP transcription factors. Additionally, our findings provide comprehensive insights into the role of the *PpGL2* gene in regulating soybean drought resistance.

## Notes

### Competing Interest Statement

The authors have declared no competing interest.

